# RIPK1 regulates β-cell fate via actions on gene expression and kinase signaling in a mouse model of β-cell self-reactivity

**DOI:** 10.1101/2025.09.18.676878

**Authors:** Christopher J. Contreras, Noyonika Mukherjee, Arianna Harris-Kawano, Egan G. Mather, Nansalmaa Amarsaikhan, Christopher Davis, Christine A. Berryhill, Madeline Peyton, Debjyoti Kundu, Kaitlyn A. Colglazier, Addison S. Miller, Renato C.S. Branco, Travis S. Johson, Steven P. Angus, Sylvaine You, Erica P. Cai, Andrew T. Templin

**Affiliations:** Department of Medicine, Roudebush VA Medical Center & Indiana University School of Medicine, Indianapolis, IN; Department of Biochemistry & Molecular Biology, Indiana University School of Medicine, Indianapolis, IN; Lilly Diabetes Center of Excellence, Indiana Biosciences Research Institute, Indianapolis, IN; Department of Pediatrics, Indiana University School of Medicine, Indianapolis, IN; Department of Biostatistics & Health Data Science, Indiana University School of Medicine, Indianapolis, IN; Université Paris Cité, Institut Cochin, CNRS, INSERM, Paris, France; Center for Diabetes & Metabolic Diseases, Indiana University School of Medicine, Indianapolis, IN

## Abstract

Type 1 diabetes (T1D) is characterized by autoimmune destruction of pancreatic β-cells, insulin insufficiency, and hyperglycemia. Receptor interacting protein kinase 1 (RIPK1) is a multifunctional regulator of cell fate with kinase and scaffolding functions, and we previously identified RIPKs as regulators of β-cell cytotoxicity *in vitro*. Here we report that Ripk1 expression is increased in islets from aged non-obese diabetic (NOD) mice and β-cells from T1D donors, suggesting that RIPK1 may drive cytokine- and autoimmune-mediated β-cell demise in T1D. Using NIT-1 β-cells derived from NOD mice, we observed that TNFα+IFNγ increase RIPK1 phosphorylation, caspase 3/7 activity, and cell death. In contrast, this cytotoxicity was blocked with small molecule RIPK1 inhibition or in Ripk1 gene-edited (Ripk1Δ) β-cells. Co-labeling of caspase 3/7 activation and cell death in single cells revealed protection from caspase-dependent and -independent forms of death in Ripk1Δ cells. RNAseq uncovered differential cell death-, immune-, and identity-related gene expression, and kinome profiling identified changes in MAPK, Eph, JAK, and other kinase activity associated with protection from cell death in RIPK1 deficient β-cells. Furthermore, *in vitro* co-culture assays and *in vivo* adoptive transfer experiments revealed that NIT-1 Ripk1Δ cells were protected from autoimmune destruction by splenocytes isolated from diabetic NOD mice. Collectively, our findings indicate that RIPK1 promotes β-cell demise in response to cytokine and autoimmune stress via actions on gene expression and kinase signaling.

Therapeutics targeting RIPK1 may provide novel opportunities for prevention or treatment of autoimmune diabetes.

## INTRODUCTION

Type 1 diabetes (T1D) is a chronic disease characterized by autoimmune killing of insulin-producing β-cells, leading to insulin insufficiency and hyperglycemia^1,2^. Interactions between β-cells and immune cells underlie β-cell death in T1D via cytokine signaling that elicits immunogenic β-cell stress^3,4^, loss of immune tolerance, and autoreactive T-cell mediated β-cell destruction^5,6^. In this context, several studies have identified functional and transcriptional β-cell stress responses that promote inflammation and immunogenicity^7–11^, including generation of neoantigens^7,8^, increased MHC class I expression^10–12^, and inflammatory cell death^13–15^. TNFα and IFNγ are proinflammatory cytokines recognized as mediators of β-cell demise in T1D. Numerous preclinical studies identified cytotoxic effects of TNFα and IFNγ on mouse and human β-cells *in vitro* and *in vivo*^11,13,16–18^. In addition, recent clinical trials found blockade of TNFα^19,20^ or JAK signaling^21^ preserves endogenous insulin secretion and improves glucose homeostasis in people with new onset T1D^19,20^, consistent with improved β-cell function and survival. However, the specific cellular and molecular mechanisms that underlie these observations remain obscure. Given our limited understanding of the complex interactions between β-cells and immune cells in T1D pathogenesis, studies to better characterize immunogenic β-cell stress and autoimmune β-cell destruction are needed to combat this disease.

Receptor interacting protein kinase 1 (RIPK1) is a multifunctional regulator of cell survival and death that acts downstream of diverse cytotoxic stimuli including death receptor ligands, interferons (IFNs), toll-like receptor (TLR) ligands, and others^22–25^. It has both kinase^22^ and scaffolding functions^26^, and these coordinate its actions on signal transduction^22,25,27^, inflammatory gene expression^27,28^, caspase-dependent apoptosis^29,30^, and caspase-independent necroptosis^31,32^, a lytic and immunogenic form of cell death. Human RIPK1 mutations are linked to inflammatory and autoimmune pathologies^33–35^. Given the potential role of RIPK1 in diabetogenic β-cell cytotoxicity, prior studies tested a small molecule RIPK1 kinase inhibitor called necrostatin-1 (Nec-1), with findings largely in agreement that RIPK1 kinase inhibition protects β-cell lines, mouse, porcine, and human islets from cytotoxicity *in vitro*^36–38^. In addition, Nec-1 prevents β-cell loss in a zebrafish model of overnutrition^39^, and treatment of porcine islets with Nec-1 prior to transplantation in diabetic mice was found to improve insulin release and glycemia *in vivo*^40^. Moreover, our earlier work demonstrated that Ripk1 gene editing blocks TNFα-induced cell death in NIT-1 β-cells derived from the non-obese diabetic (NOD) mouse model of T1D^14^. In contrast, mice harboring Ripk1^S25D/S25D^ or Ripk1^D138N/D138N^ mutations that reduce RIPK1 kinase activity were not protected from hyperglycemia following streptozotocin (STZ) treatment, a model of chemically-induced diabetes^41,42^. Although RIPK1 has emerged as a stress sensor with effects on inflammation, immune regulation, and programmed cell death^23,25,29,43^, and despite mounting evidence RIPK1 is a regulator of β-cell survival, few studies have characterized its impact on β-cell demise in autoimmune diabetes.

We hypothesized that RIPK1 regulates cytokine- and autoimmune-mediated β-cell demise in T1D. To test this hypothesis, we used control (CTL) and Ripk1 gene-edited (Ripk1Δ) NIT-1 β-cells derived from the NOD mouse model of spontaneous β-cell self-reactivity. We exposed these cells to TNFα and IFNγ or co-cultured them with self-reactive splenocytes isolated from diabetic NOD mice. We performed high content, automated live cell imaging and analysis, RNA sequencing, multiplex kinase inhibitor bead affinity chromatography-mass spectrometry (kinome profiling), and flow cytometry, and we tested a small molecule RIPK1 kinase inhibitor (SZM’679^44^) and protein degrader (LD4172^45^). We also evaluated the role of β-cell RIPK1 in self-reactivity and autoimmune killing *in vivo* using an NOD mouse β-cell implantation model. This study provides evidence that RIPK1 is a T1D-relevant mediator of β-cell signal transduction, cytotoxicity, and autoimmunity.

## RESULTS

### β-cell RIPK1 is activated by proinflammatory cytokines, and its expression is increased in aged NOD islets and β-cells from humans with T1D

RIPK1 is a multifunctional protein with kinase and scaffolding domains (RHIM, death) that coordinately regulate its actions on cell fate (**Fig. 1A**). Given that RIPK1 is a mediator of TNFα and IFNγ signaling in other cell types^24,26,29,46^, we evaluated the effects of TNFα+IFNγ treatment on RIPK1 expression and phosphorylation in NIT-1 mouse β-cells (**Fig. 1B**), mouse islets (**Fig. 1C**), and EndoC-βH1 human β-cells (**Fig. 1D**). In each case, we found TNFα+IFNγ treatment increased Ripk1 RNA expression and RIPK1 protein phosphorylation at serine 166, indicating activation of RIPK1 under these conditions^47^ (**Fig. 1B-D**). To characterize RIPK1 expression in islets, we performed immunohistochemistry for RIPK1 and insulin in pancreas sections from non-diabetic mouse and human donors. In both species, RIPK1 immunoreactivity was abundant in islets and was largely colocalized with insulin, indicating β-cells express RIPK1 protein *in situ* (**Fig. 1E,G**). We also found that Ripk1 RNA expression is significantly increased in NOD mouse islets as they age from 8 to 20 weeks (**Fig. 1F**), a period during which β-cell autoimmunity and cytotoxicity progress *in vivo*. Drawing from the Human Pancreas Analysis Program (HPAP) single cell transcriptomics data set^48,49^, we observed that RIPK1 RNA expression is elevated in β-cells from T1D versus non-diabetic donors (**Fig. 1H**). These findings suggest that RIPK1 is a T1D-relevant kinase and led us to undertake additional studies to decipher the role of RIPK1 in cytokine- and autoimmune-mediated β-cell demise.

**Figure 1:**
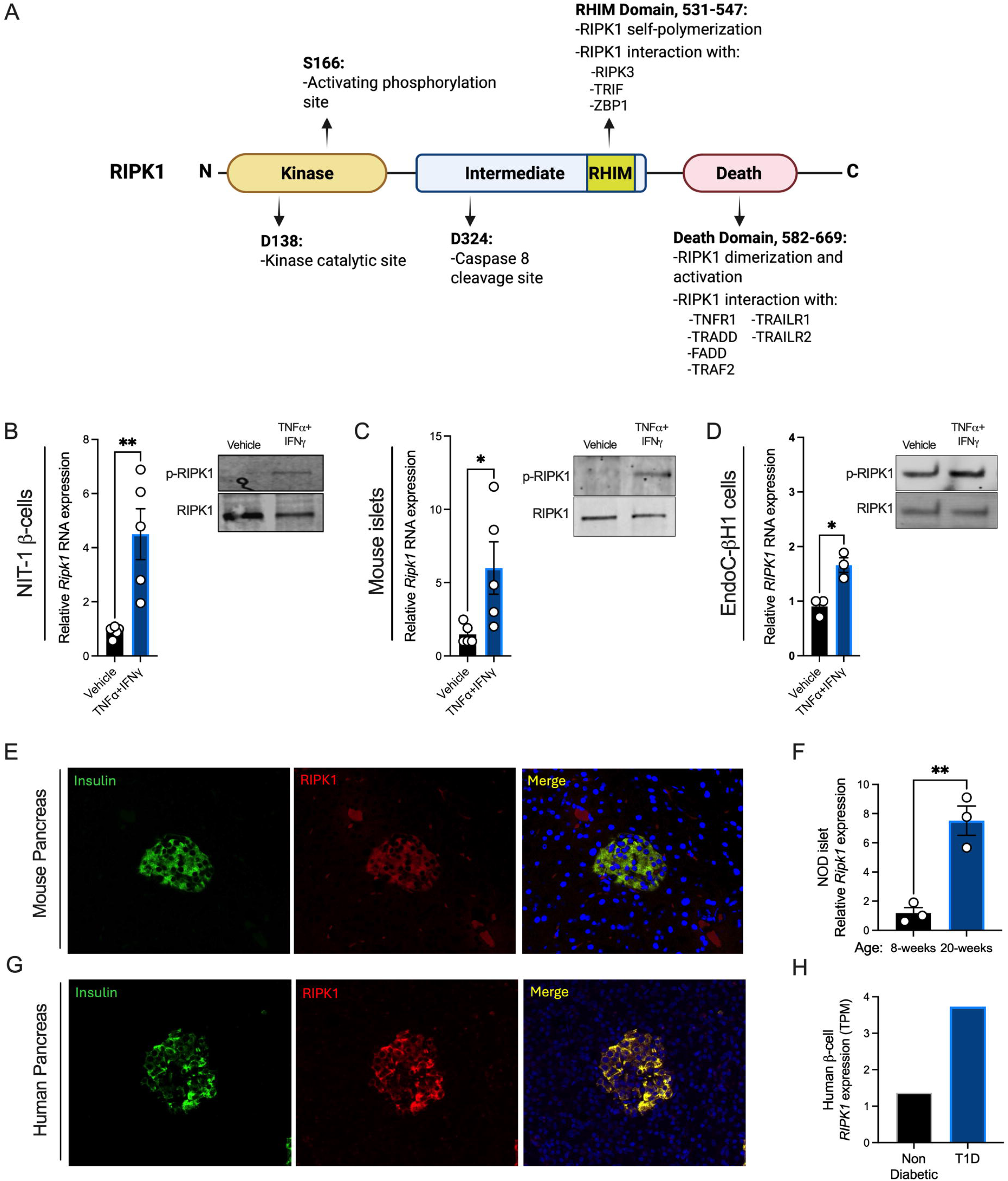
β-cell RIPK1 is activated by proinflammatory cytokines, and its expression is increased in aged NOD islets and β-cells from humans with T1D. **A)** Schematic of receptor interacting protein kinase 1 (RIPK1) amino acid residues, domains, and functions. **B)** Vehicle or TNFα (40 ng/mL) + IFNγ (100 ng/mL) were applied to mouse NIT-1 β-cells (4 hours), **C)** mouse islets (18 hours), and **D)** human EndoC-βH1 cells (18 hours), then Ripk1 RNA expression was quantified by qPCR (left) and RIPK1 phosphorylation was visualized by immunoblot (right). **E)** Mouse and **G)** human pancreas sections were stained for insulin (green) and RIPK1 (red) to evaluate RIPK1 expression in beta cells *in situ*. **F)** Ripk1 RNA expression was quantified in islets from NOD mice at 8- and 20-weeks of age. **H)** HPAP single cell RNA expression data was queried for RIPK1 expression in β-cells from human donors without or with T1D. Data are represented as mean ± SEM. n=3-5. *p < 0.05; **p < 0.01

### Small molecule RIPK1 kinase inhibition or protein degradation protects NIT-1 β-cells from cytokine-induced death

We tested two recently reported RIPK1-targeting small molecules to evaluate their effects on TNFα+IFNγ-induced β-cell death. To compare the effects of RIPK1 kinase inhibition versus protein degradation, we evaluated a small molecule RIPK1 kinase inhibitor (SZM’679^44^) and a RIPK1-directed proteolysis targeting chimera (PROTAC, LD4172^45^) (**Fig. 2A, B**). We first validated the effectiveness of LD4172 to diminish RIPK1 protein expression in NIT-1 CTL cells. We found that LD4172 reduced RIPK1 protein abundance by ∼30% after 2 hours, and this reduction reached >75% after 24 hours (**Fig. 2C**). We next quantified cell death using an Incucyte S3 live cell imaging and analysis system and Sytox green, a membrane impermeable DNA binding dye^50–52^. Neither SZM’679 nor LD4172 treatment alone altered cell death over 24 hours (**Fig. 2D, E**). In contrast, TNFα+IFNγ treatment increased NIT-1 cell death over this interval, and co-treatment with either SZM’679 or LD4172 diminished TNFα+IFNγ-induced NIT-1 cell death (**Fig. 2D, E**). The effects of SZM’679 and LD4172 on caspase 3/7 activity were assessed using a static luminometric caspase 3/7 assay. Following pretreatment with either SZM’679 or LD4172, NIT-1 cells were treated with TNFα+IFNγ for an additional 4 hours, then caspase 3/7 activity was quantified. RIPK1 kinase inhibition with SZM’679 effectively blocked cytokine-induced caspase 3/7 activation, whereas RIPK1 protein degradation with LD4172 did not have this effect (**Fig. 2F**). These studies indicate that small molecules targeting RIPK1 can counteract TNFα+IFNγ-induced β-cell cytotoxicity.

**Figure 2:**
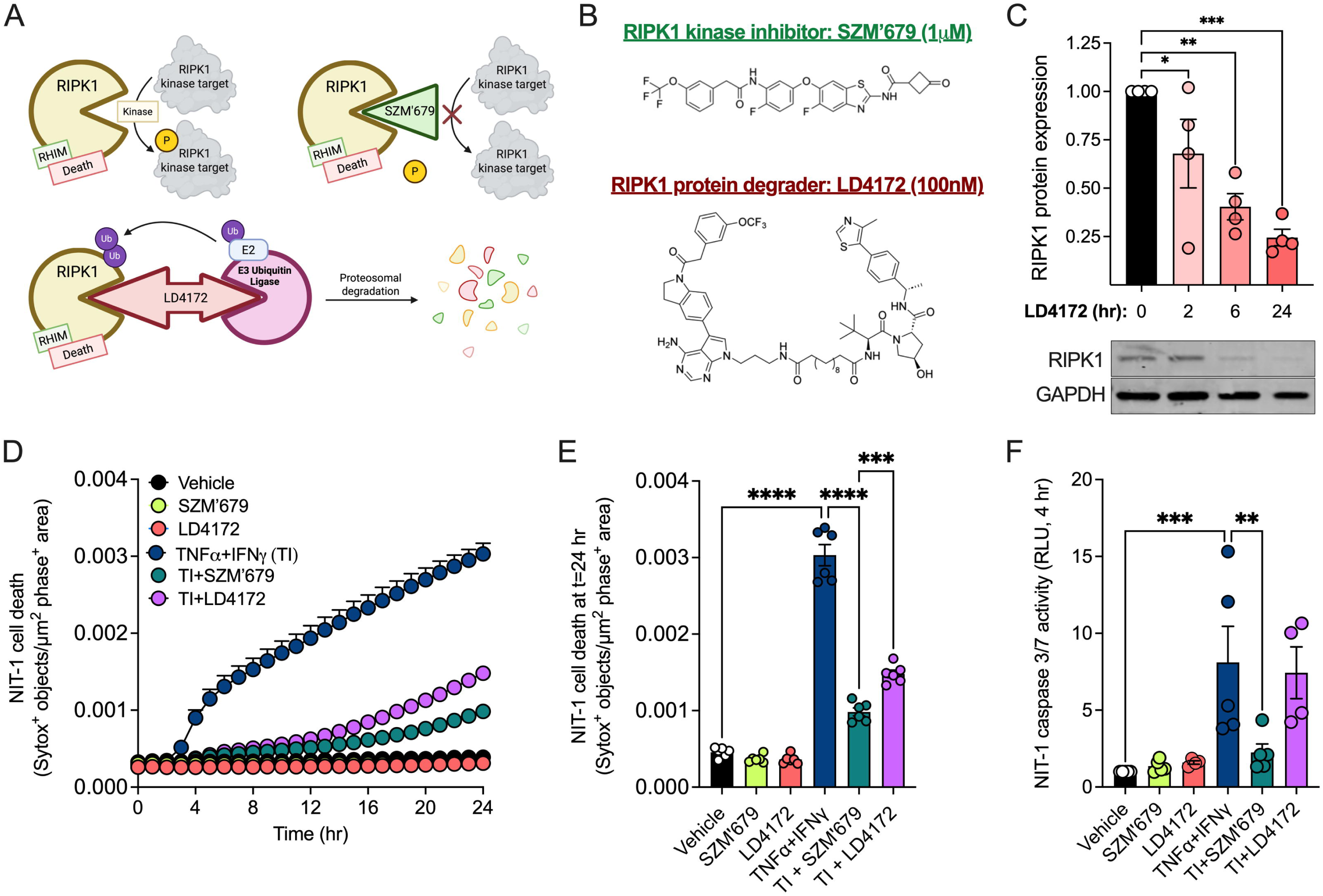
Small molecule RIPK1 kinase inhibition or protein degradation protects NIT-1 β-cells from cytokine-induced death. **A)** Effects of small molecule inhibitors SZM’679 (kinase inhibitor, green) and LD4172 (PROTAC, red) on RIPK1. **B)** Chemical structures of SZM’679 and LD4172. **C)** NIT-1 cell RIPK1 protein expression was quantified by immunoblot 0, 2, 6, and 24 hours after treatment with LD4172. NIT-1 cells were treated with vehicle, SZM’679 (1 μM), LD4172 (100 nM), TNFα (40 ng/mL) + IFNγ (100 ng/mL) (TI), TI + SZM’679, or TI + LD4172. SZM’679 and LD4172 were added 4 hours prior to treatment with TNFα+IFNγ. **D)** Cell death was measured as Sytox green positive objects per μm^2^ phase positive area hourly for 24 hours, and **E)** cell death was quantified at 24 hours. **F)** Caspase 3/7 activity was quantified in NIT-1 cells using a luminometric caspase 3/7 assay. Data are represented as mean ± SEM. n=4-6. *p < 0.05; **p < 0.01; ***p < 0.001, ****p < 0.0001

### Ripk1 gene editing protects NIT-1 β-cells from caspase-dependent and caspase-independent death following cytokine treatment *in vitro*

To further interrogate the role of RIPK1 in β-cell cytotoxicity, we utilized a genetic engineering approach to modify the Ripk1 gene in β-cells. NIT-1 β-cells were subjected to CRISPR-Cas9 gene editing with either guide RNAs targeting exons 2 and 3 of Ripk1 (NIT-1 Ripk1Δ) or scrambled control guide RNA (NIT-1 CTL), resulting in generation of two NIT-1 β-cell populations. Ripk1 gene editing was confirmed in NIT-1 Ripk1Δ cells by genomic DNA PCR^50^, and immunoblot analysis showed a 65% reduction in RIPK1 protein expression in the NIT-1 Ripk1Δ cell line (**Fig. 3A**). We next conducted real time cell death assays using Sytox green to quantify death in NIT-1 CTL and NIT-1 Ripk1Δ cells (**Fig. 3B,F**). As expected, NIT-1 CTL cells were highly susceptible to TNFα+IFNγ-induced cell death over 24 hours. In contrast, NIT-1 Ripk1Δ cells displayed significant protection from cell death under these conditions (**Figs. 3C,D**). Under vehicle treatment, cell death was not different between NIT-1 CTL versus Ripk1Δ cells at any time (**Figs. 3C,D**). Static quantification of caspase 3/7 activation after TNFα+IFNγ exposure demonstrated that caspase 3/7 activity was increased nearly 6-fold in NIT-1 CTL after 4 hours, while caspase activation was absent in NIT-1 Ripk1Δ cells (**Fig. 3E**). Like our findings with small molecule RIPK1 inhibitors, these data indicate that RIPK1 mediates cytokine-induced caspase 3/7 activation and cell death in NIT-1 β-cells.

**Figure 3:**
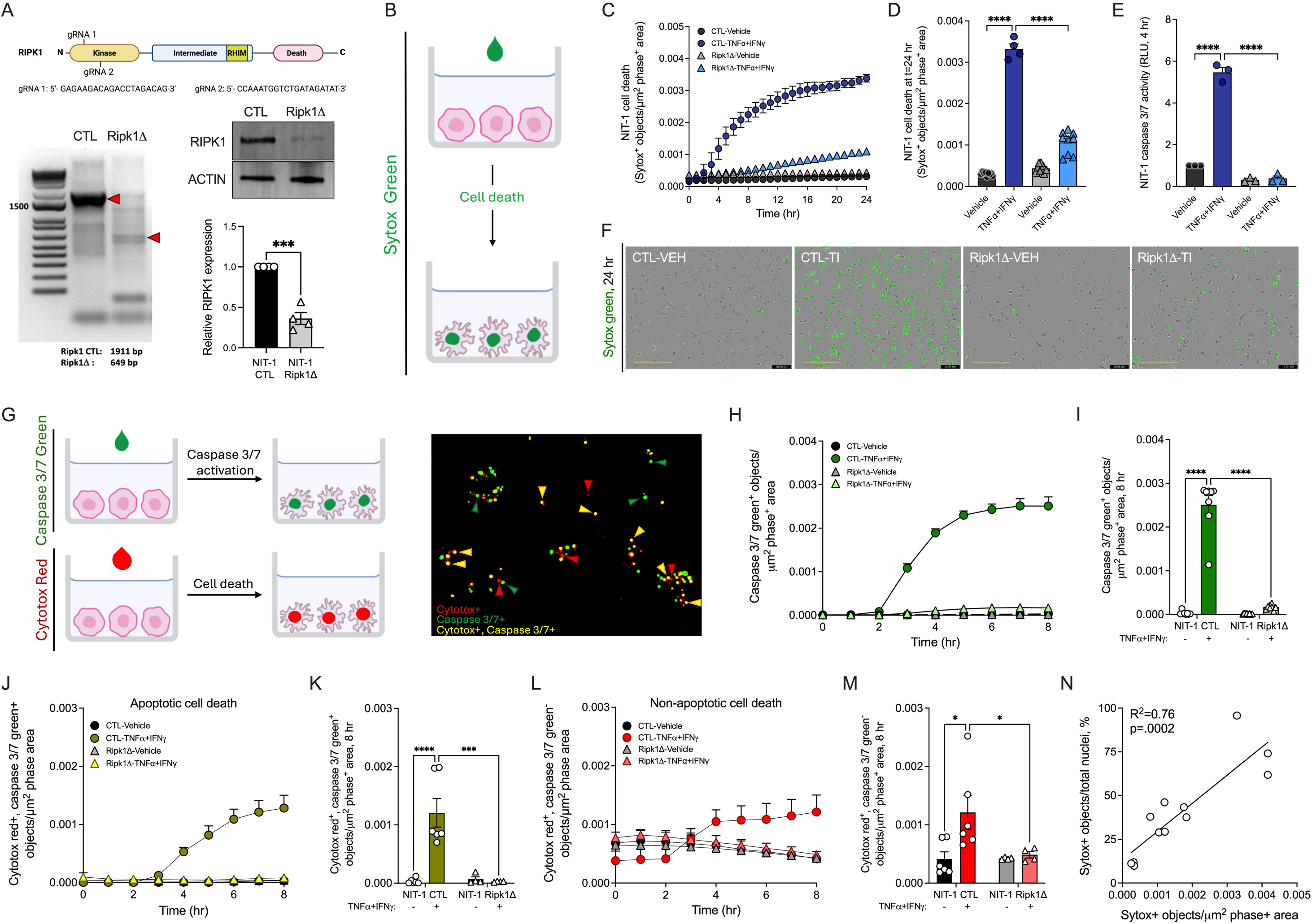
Ripk1 gene editing protects NIT-1 β-cells from caspase-dependent and caspase-independent death following cytokine treatment *in vitro*. **A)** Illustration of Ripk1 gene targeting strategy used to generate NIT-1 Ripk1Δ cells (top). PCR of Ripk1 genomic DNA was performed in NIT-1 CTL versus Ripk1Δ cells (bottom left). RIPK1 protein expression was quantified in NIT-1 CTL versus Ripk1Δ cells via immunoblot analysis, and relative RIPK1 protein expression was quantified (bottom right). **B)** Sytox green was used to label dying cells following membrane permeabilization. **C)** Cell death was measured as Sytox green positive objects per µm^2^ phase positive area in NIT-1 CTL versus NIT-1 Ripk1Δ cells treated with vehicle or TNFα (40 ng/mL) + IFNγ (100 ng/mL) hourly for 24 hours, **D)** cell death was quantified at 24 hours, and **F)** representative images are shown. **E)** Caspase 3/7 activity was quantified in NIT-1 CTL and Ripk1Δ cells 4 hours post treatment. **G)** Caspase 3/7 green and Cytotox red were used in tandem to identify single cells undergoing caspase 3/7 activation and/or cell death. **H-M)** NIT-1 CTL and NIT-1 Ripk1Δ cells were treated with vehicle or TNFα+IFNγ, then **H,I)** total Caspase 3/7 green positive objects (green), **J,K)** Cytotox red, Caspase 3/7 green dual-positive objects (yellow), and **L,M)** Cytotox red positive, Caspase 3/7 green negative objects (red) were quantified over 8 hours. **N)** Correlation of Sytox green^+^/μm^2^ phase^+^ area values and Sytox green^+^ objects/Nuclight red^+^ object values (%) was performed. Data are represented as mean ± SEM. n=3-9. *p < 0.05; *** p < 0.001; ****p < 0.0001

To further investigate the relationship between caspase 3/7 activation and cell death in TNFα+IFNγ-induced β-cell cytotoxicity, we used Cytotox red (a membrane impermeable DNA binding dye that labels dying cells) and Caspase 3/7 green (a membrane permeable caspase 3/7 substrate that binds DNA following cleavage by caspase 3 or 7) (**Fig. 3G**, left). This live cell imaging approach allowed us to quantify both caspase 3/7 activation and cell death in single cells in real time (**Fig. 3G**, right). Using this approach, we found that caspase 3/7 activity was increased 2-3 hours post TNFα+IFNγ-treatment in NIT-1 CTL cells and reached maximal activity 8 hours post treatment (**Fig. 3H,I**). In contrast, TNFα+IFNγ-treatment failed to increase caspase 3/7 activity in NIT-1 Ripk1Δ cells over this period (**Fig. 3H,I**). We next quantified apoptotic (defined as double Cytotox red^+^, caspase 3/7 green^+^ objects) and non-apoptotic (defined as Cytotox red^+^, caspase 3/7 green^-^ objects) NIT-1 β-cells following treatment with TNFα+IFNγ. We found NIT-1 CTL cells undergo both caspase-dependent (**Figs. 3J,K**) and caspase-independent (**Figs. 3L,M**) forms of cell death in response to TNFα+IFNγ treatment over 8 hours, and both forms of cell death were blocked in NIT-1 Ripk1Δ cells (**Figs. 3J-M**). To relate our cell death metric (fluorescent^+^ objects per μm^2^ phase^+^ area) to percent cell death, we utilized Nuclight red, a membrane permeable DNA binding dye that labels all nuclei. We then compared Sytox green^+^/μm^2^ phase^+^ area values with Sytox green^+^ objects/Nuclight red^+^ object values (%) within wells to correlate our cell death units to percent cell death (**Fig. 3N**).

### Protection from cytokine-induced cell death is associated with altered gene expression in Ripk1 deficient NIT-1 β-cells

Although cytotoxic functions of RIPK1 are generally attributed to its phosphorylation-mediated signal transduction functions, recent studies indicate RIPK1 can mediate inflammatory gene expression^22,27,28^. To characterize the role of RIPK1 in NIT-1 β-cell gene expression under cytokine stress, we performed RNA sequencing on NIT-1 CTL and Ripk1Δ cells treated with vehicle or TNFα+IFNγ. This approach allowed us to test effects of RIPK1 independent of treatment (**Figs. 4A,B**), effects of TNFα and IFNγ independent of genotype (**Figs. 4C,D**), and interaction effects where the treatment response differed between genotypes (**Figs. 4E,F**). For each comparison, we visualized the top 10 most significantly upregulated and downregulated genes by FDR (volcano plots, **Figs. 4A,C,E**). We also employed a combined ranking approach to integrate statistical significance and magnitude of fold change to identify genes of interest (heatmaps, **Figs. 4B,C,F**).

**Figure 4:**
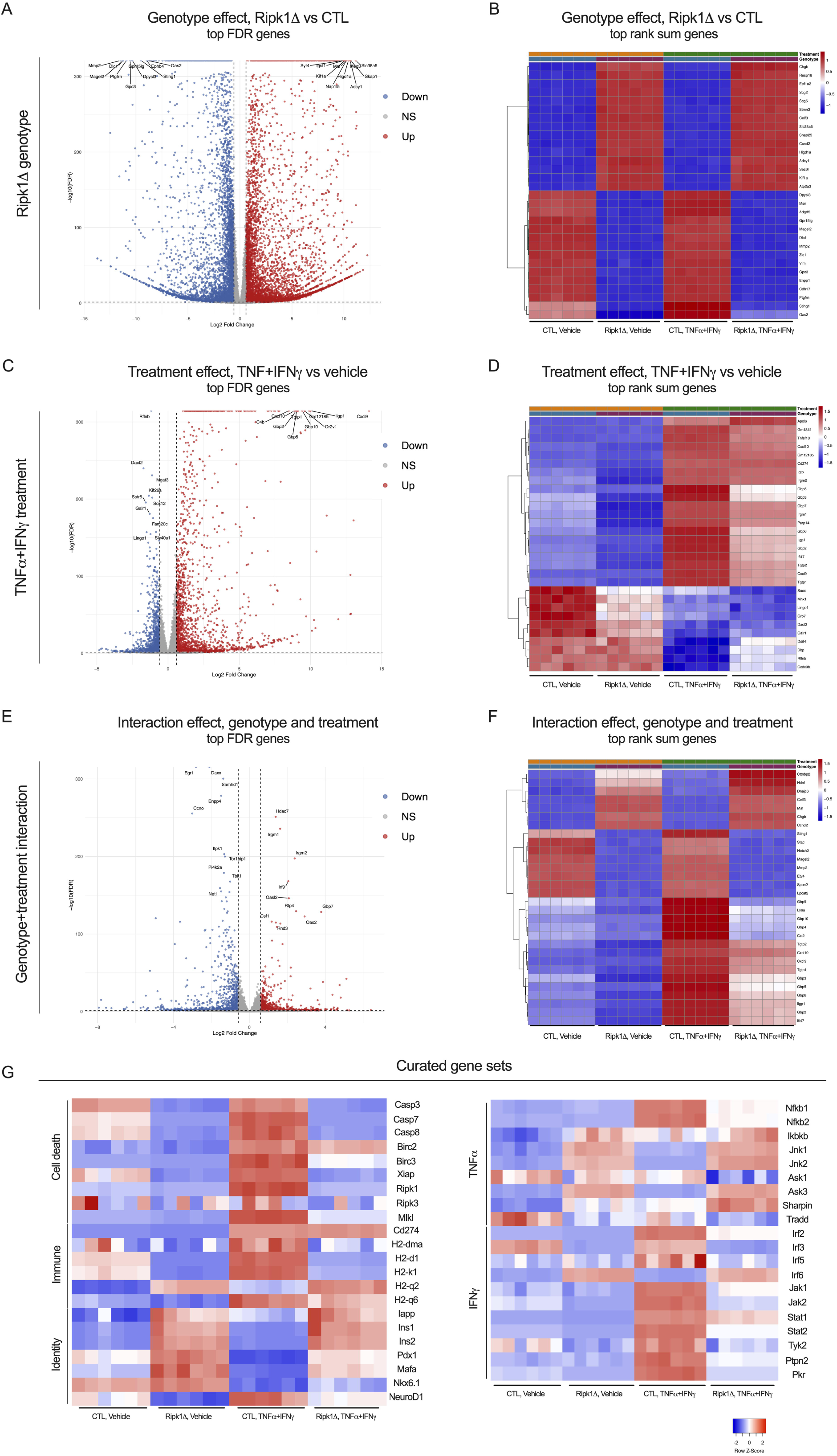
Protection from cytokine-induced cell death is associated with altered gene expression in Ripk1 deficient NIT-1 β-cells. NIT-1 CTL and Ripk1Δ cells were treated with vehicle or TNFα (40 ng/mL) + IFNγ (100 ng/mL) for 4 hours, then RNA was isolated and bulk RNA sequencing was performed. Following quality control analyses, RNA sequencing datasets were examined to test effects of **A,B)** Ripk1 genotype independent of treatment, **C,D)** effects of TNFα and IFNγ independent of genotype, and **E,F)** effects of interaction between genotype and treatment. Red, upregulated gene expression; blue, downregulated gene expression. **A,C,E)** For each comparison, the top 10 most significantly upregulated and downregulated genes by FDR were visualized by volcano plot. **B,D,F)** The top 30 genes in each comparison were identified using a combined ranking approach that integrates statistical significance and magnitude fold change, and these were visualized using heatmaps. **G)** Curated heatmaps depict cell death-, immune-, cytokine-, and identity-related genes that are differentially expressed in NIT-1 Ripk1Δ versus CTL cells. n=6.

Our results revealed clear differences in gene expression between NIT-1 CTL and Ripk1Δ cells, including upregulation of *Atp2a3/Serca3* and *Syt4*, genes involved in calcium homeostasis and insulin secretion, and downregulation of *Sting1* and *Oas2*, genes involved in innate immune response and interferon signaling, among many others (**Figs. 4A,B**). Our studies also revealed many genes that were upregulated by TNFα+IFNγ treatment independent of genotype including *Cxcl9*, *Cxcl10*, *Irgm1*, and *Irgm2*, genes involved in chemotaxis, innate immune response, and interferon signaling. Genes downregulated by TNFα+IFNγ treatment independent of genotype such as *Rflnb*, *Mnx1*, *Dact2*, and *Grb7* were also identified (**Figs. 4C,D**). Notably, there were several genes whose expression in response to TNFα+IFNγ exposure differed in NIT-1 Ripk1Δ compared to NIT-1 CTL cells, and we annotated these genes as displaying an interaction effect. Genes identified as being more highly upregulated in Ripk1Δ versus CTL cells following treatment include *Irgm1*, *Irgm2*, *Irf9, Oas2,* and *Oasl2,* and genes more highly downregulated in Ripk1Δ versus CTL under cytokine treatment include *Daxx*, *Tbk1*, *Ccl2*, *Gbp4*, and *Gbp9* (**Figs. 4E,F**). We also visualized expression of a curated set of genes related to cell death, immune response, and β-cell identity, as well as TNFα and IFNγ signaling (**Fig. 4G**). Several cytotoxicity-related genes including *Casp3*, *Casp7*, *Mlkl*, *Nfkb1*, *Nfkb2*, *Jnk1*, *Jnk2*, *Jak1*, *Jak2*, *Stat1*, *Stat2,* and *Tyk2* were found to exhibit RIPK1-related expression patterns (**Fig. 4G**). In sum, these findings reveal significant effects of RIPK1 on NIT-1 β-cell gene expression that are relevant to cytokine and autoimmune stress.

### Protection from cytokine-induced cell death is associated with altered kinase signaling in Ripk1 deficient NIT-1 β-cells

To test the role of RIPK1 in NIT-1 β-cell kinase signaling under cytokine-induced stress conditions, we performed kinome profiling (multiplex kinase inhibitor bead affinity chromatography-mass spectrometry, MIB-MS^53^) on lysates from NIT-1 CTL and Ripk1Δ cells treated with vehicle or TNFα+IFNγ (**Fig. 5A**). This chemical-proteomics method utilizes kinase inhibitor-conjugated beads to capture active kinases which are identified by mass spectrometry. To our knowledge, these studies represent the first use of this approach to quantify the functional kinome in β-cells. Our experiments identified kinases whose capture was altered with loss of RIPK1 under vehicle treatment (**Figs. 5B,D**) and/or TNFα+IFNγ treatment conditions (**Figs. 5C,E**), and these changes were visualized with volcano plots (**Figs. 5B,C**) and heatmaps (**Figs 5. D,E**). Venn diagrams illustrate kinases whose activity was down- or up-regulated in NIT-1 Ripk1Δ cells under vehicle treatment, cytokine treatment, or both (**Fig. 5F,G**). We confirmed that RIPK1 kinase binding was reproducibly downregulated in NIT-1 Ripk1Δ cells (p=0.055). We also identified prominent changes in MAPK, Eph, JAK, and other kinase signaling cascades in RIPK1 deficient β-cells. To identify β-cell kinase signaling pathways regulated by RIPK1, we performed Gene Ontology (GO) enrichment analysis on lists of kinases that were downregulated (**Fig. 5H**) or upregulated (**Fig. 5I**) in NIT-1 Ripk1Δ cells. This analysis yielded several RIPK1-regulated processes, including MAPK signaling, JNK signaling, cell death, ephrin receptor activity, and ubiquitin protein ligase binding (**Figs. 5H,I**). These findings warrant further investigation and suggest that RIPK1 is a key regulator of cytotoxicity-relevant kinase activity in β-cells.

**Figure 5:**
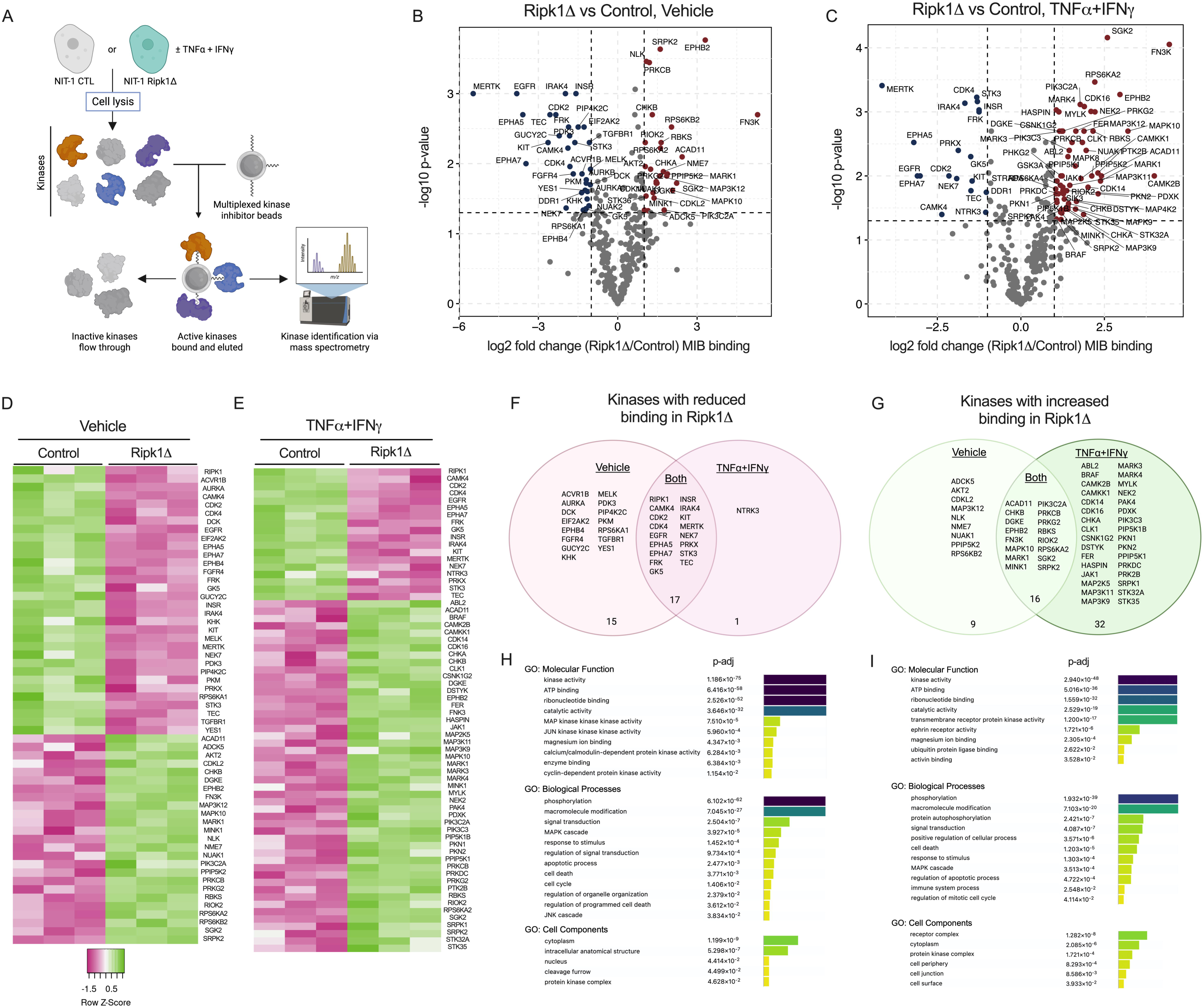
Protection from cytokine-induced cell death is associated with altered kinase signaling in Ripk1 deficient NIT-1 β-cells. **A)** Schematic for kinome profiling/MIB-MS workflow in NIT-1 CTL and Ripk1Δ cells treated with vehicle or TNFα (40 ng/mL) + IFNγ (100 ng/mL) for 4 hours. Volcano plots of differentially activated kinases in NIT-1 Ripk1Δ versus CTL cells under **B)** vehicle, and **C)** TNFα+IFNγ treatment conditions. Red, upregulated kinase activity expression; blue, downregulated kinase activity. Heatmaps illustrate differential kinase activity under **D)** vehicle, and **E)** TNFα+IFNγ treatment conditions. Z-scores; green, high kinase binding; magenta, low kinase binding. Venn diagrams illustrate kinases that are F) downregulated or G) upregulated under vehicle treatment, TNFα+IFNγ treatment, or both. Gene ontology (GO) enrichment analysis identified significantly enriched processes from kinases that are **H)** downregulated or **I)** upregulated in NIT-1 Ripk1Δ cells. n=3.

### Ripk1 deficient NIT-1 β-cells are protected from autoimmune-mediated killing *in vitro* and *in vivo*

To query the role of RIPK1 in β-cell directed immune responses in autoimmune diabetes, we isolated splenocytes from diabetic NOD mice and co-cultured them with either NIT-1 CTL or Ripk1Δ cells *in vitro* (**Fig. 6A**). ELISPOT assays showed that splenocyte IFNγ production increased after stimulation with NIT-1 β-cells, and this IFNγ production was equivalent in NIT-1 CTL and Ripk1Δ co-cultures (**Fig. 6B,C**). Similarly, flow cytometry revealed comparable degrees of CD69 expression on CD8^+^ T-cells co-cultured with NIT-1 CTL or Ripk1Δ β-cells (**Fig. 6D**), suggesting T-cell activation is not altered by β-cell RIPK1 expression. In contrast, diabetic splenocyte-mediated β-cell death was halved in NIT-1 Ripk1Δ versus CTL cells following splenocyte co-culture (**Fig. 6E**), indicative of a β-cell intrinsic protection from autoimmune-killing with loss of RIPK1 *in vitro*.

**Figure 6:**
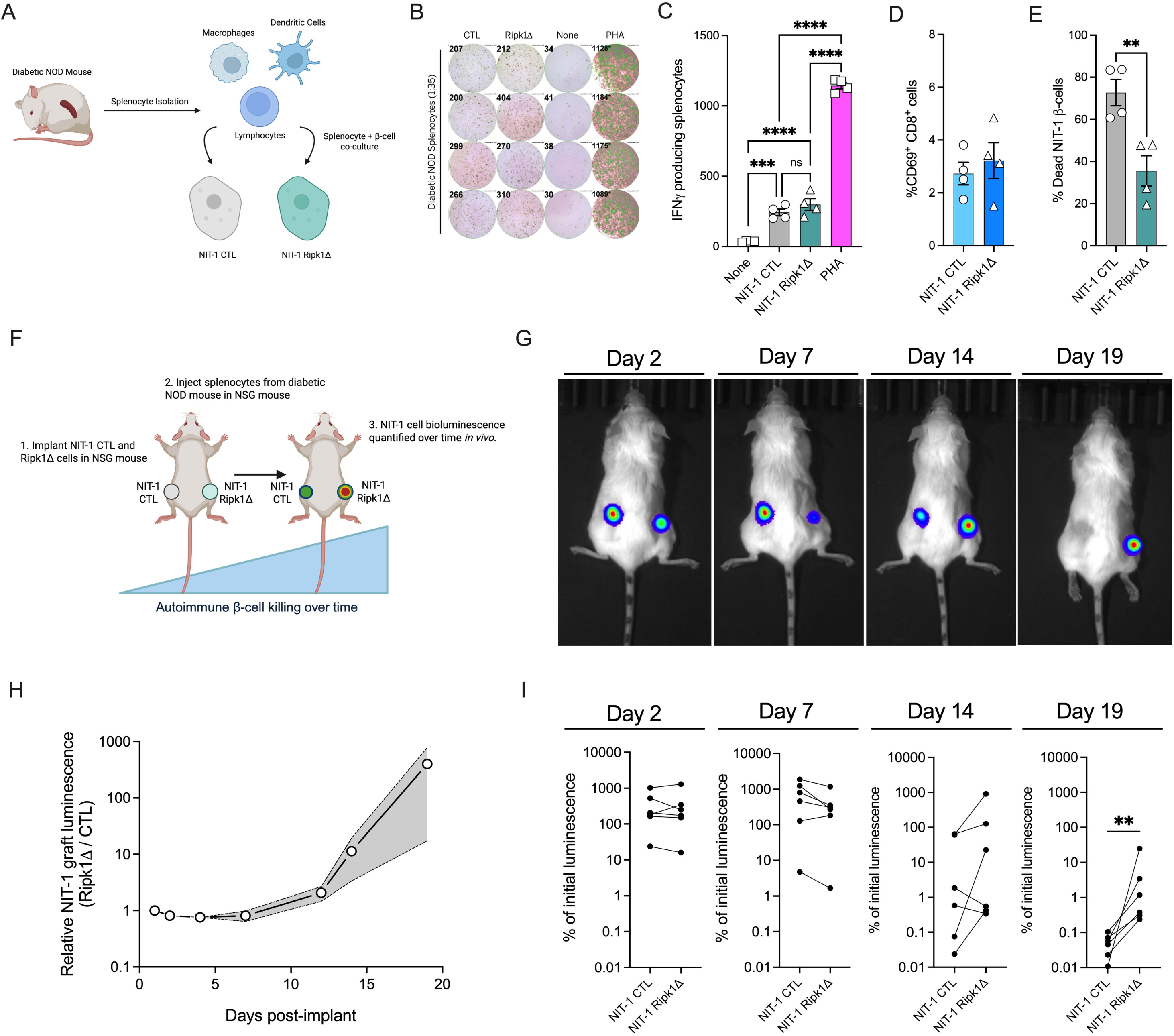
Ripk1 deficient NIT-1 β-cells are protected from autoimmune-mediated killing *in vitro* and *in vivo*. **A)** Splenocytes isolated from spontaneously diabetic NOD mice were co-cultured with NIT-1 CTL or Ripk1Δ cells. **B)** IFNγ production was quantified following β-cell and splenocyte co-culture (1:35 ratio) for 24 hours. Splenocytes alone (none) served as negative controls and phytohemagglutinin (PHA) treated splenocytes served as positive controls. **C)** IFNγ producing splenocytes were quantified via ELISPOT assay. **D,E)** NIT-1 cells were co-cultured with splenocytes (1:2 ratio) for 48 hours, then flow cytometry was performed to quantify **D)** CD69+CD8+ T-cell abundance and **E)** % NIT-1 cell death (CTV+Zombie+). **F)** Schematic of *in vivo* autoimmune NIT-1 cell killing assay. NIT-1 CTL and Ripk1Δ cell graft luminescence was visualized at days 1, 2, 4, 7, 12, 14, and 19 post implant. **G)** Representative images of IVIS graft imaging 2, 7, 14, and 19 days post implant. **H)** Luminescence of NIT-1 Ripk1Δ cell grafts relative to NIT-1 CTL cell grafts was quantified over the 19 day study period. **I)** Luminescence of NIT-1 CTL and Ripk1Δ cell grafts 2, 7, 14, and 19 days post implant represented as % of graft luminescence at day 1. Each line represents a NIT-1 CTL cell graft (left) and a NIT-1 Ripk1Δ cell graft (right) from a single mouse. n=4-6.

We next undertook studies to assess the role of RIPK1 in autoimmune-mediated β-cell killing *in vivo*. Here, we implanted luciferase-expressing NIT-1 CTL and NIT-1 Ripk1Δ cells on opposite flanks of immunodeficient NOD.scid.gamma (NSG) mice, induced β-cell autoimmunity via intravenous injection of splenocytes isolated from spontaneously diabetic NOD mice, then quantified NIT-1 graft luminescence 1, 2, 4, 7, 12, 14, and 19 days post-implant using an IVIS *in vivo* imaging system (**Fig. 6F,G**). NIT-1 CTL and Ripk1Δ cell graft luminescence was similar following implantation and remained unchanged over the initial 7-day study period (**Fig. 6H,I**). However, by day 14 post-implant, NIT-1 Ripk1Δ cell graft luminescence was 11-fold greater than in NIT-1 CTL grafts, and this difference increased to >100-fold Ripk1Δ/CTL graft luminescence by day 19 post-implant (**Fig 6H**). At day 19, NIT-1 CTL grafts averaged 0.05% of initial graft luminescence, whereas NIT-1 Ripk1Δ grafts averaged 5.1% (**Fig. 6I**). These findings indicate that RIPK1 deficiency protects β-cells from autoimmune-mediated destruction via actions on β-cell survival and death signaling.

## DISCUSSION

Cytokine stress contributes to β-cell cytotoxicity in T1D in part by increasing inflammatory gene expression^54^, chemokine production^10,55^, and autoantigen presentation^11,12^, thereby stimulating immune responses against β-cells^56^. TNFα and IFNγ are recognized mediators of β-cell demise in early T1D, with preclinical studies showing TNFα and IFNγ have cytotoxic and immunogenic effects on mouse and human β-cells *in vitro* and *in vivo*^11,13,16–18^. In addition, recent clinical trials demonstrated that blockade of TNFα^19,20^ or IFN signaling^21^ preserves endogenous insulin secretion and improves glucose homeostasis in people with new onset T1D^19,20^. Although receptor interacting protein kinase 1 (RIPK1) is an essential mediator of TNFα and IFNγ signaling in non-islet cell types^25,29,43^, it has not been well studied in autoimmune diabetes. Here, we characterized the role of RIPK1 in regulation of β-cell fate under cytokine and autoimmune stress conditions using NIT-1 β-cells and autoreactive splenocytes isolated from the NOD mouse model of spontaneous autoimmune diabetes. We demonstrated that RIPK1 regulates cytokine- and autoimmune-mediated β-cell death via actions on gene expression and kinase signaling.

Our studies found that either Ripk1 gene editing or small molecule inhibition of RIPK1 blocks TNFα+IFNγ-induced NIT-1 β-cell death. Using an automated live cell imaging and analysis technique that allows quantification of caspase 3/7 activation and cell death in single cells, we noted that caspase-independent cell death increased 3 hours post TNFα+IFNγ treatment in NIT-1 CTL cells, whereas caspase-dependent cell death was not detected until 4 hours post treatment. However, the preponderance of NIT-1 cell death occurred after 4 hours in association with caspase 3/7 activation. These findings indicate apoptosis is the predominant form of NIT-1 cell death in response to TNFα+IFNγ, and that a subpopulation of NIT-1 β-cells undergo caspase 3/7 independent cell death, consistent with necroptosis^31^. RIPK1 deficient NIT-1 Ripk1Δ cells were protected from both caspase-dependent and caspase-independent forms of TNFα+IFNγ-induced cell death, indicating RIPK1 acts upstream of both cell death pathways in this model.

We utilized unbiased approaches to characterize the effects of RIPK1 on β-cell gene expression and kinase signaling, and these studies revealed stark differences in RIPK1-deficient versus control NIT-1 cells. RNAseq confirmed expected effects of RIPK1 on NFkB-related gene expression^57^ and revealed RIPK1-related expression of many other cell death-, immune-, and cytokine-related genes.

Unexpectedly, these studies also uncovered a role for RIPK1 in β-cell identity, with *Ins1*, *Ins2*, *Iapp*, and *Pdx1* among the genes upregulated in NIT-1 Ripk1Δ cells. These findings suggest a link between β-cell fate and identity and provide a rationale for future studies on the role of RIPK1 in β-cell maturity and function. To understand how RIPK1 deficiency alters kinase activity in β-cells, we utilized a novel chemical-proteomics approach called kinome profiling (described in **Fig. 5A**). These studies confirmed that RIPK1 kinase abundance is reduced in NIT-1 Ripk1Δ cells, revealed roles for RIPK1 in MAPK and JAK signal transduction under cytokine treatment, and identified several other differentially upregulated kinases including Eph receptor kinases, PKR, and MERTK, a TAM family kinase recently linked to regulation of necroptosis^58^. Interestingly, many kinases exhibited greater changes in MIB binding in NIT-1 Ripk1Δ cells than RIPK1 itself. This suggests that RIPK1 has significant effects on kinase signaling independent of its own kinase activity, and this effect may be related to its scaffolding functions. In addition, most differentially bound kinases in NIT-1 Ripk1Δ cells under TNFα+IFNγ treatment were upregulated, indicating RIPK1 both positively and negatively regulates kinase signaling networks in β-cells.

The work presented here is among the first to examine RIPK1 in mouse models of β-cell self-reactivity and autoimmunity. Our findings in NIT-1 β-cells demonstrate that RIPK1-deficiency protects from autoimmune-mediated death following exposure to self-reactive splenocytes *in vitro* and *in vivo*. We chose to use NIT-1 β-cells in these studies because they are derived from the NOD mouse. Thus, we were able to test the role of RIPK1 in β-cell autoimmunity using both β-cells and splenocytes taken from this widely used model of T1D. In addition, we previously found that NIT-1 β-cells exhibit increased susceptibility to TNFα-induced cell death compared to INS-1 β-cells^50^, suggesting potential differences in cytokine- and RIPK1-signaling in T1D-prone versus T1D-resistant β-cell lines that would be reflected in our observations. Previous reports showed that mice harboring Ripk1^S25D/S25D^ or Ripk1^D138N/D138N^ mutations that reduce RIPK1 kinase activity were not protected from hyperglycemia following STZ treatment^41,42^, a model of chemically-induced diabetes. Our unpublished findings in Ripk1^D138N/D138N^ mice are in line with these results (*data not shown*). Taken together, our observations point to a role for RIPK1 in cytokine- and autoimmune-mediated β-cell demise in spontaneous autoimmune diabetes that is not recapitulated in models of STZ-induced hyperglycemia.

Prior studies of β-cell RIPK1 have utilized necrostatin-1 (Nec-1), a small molecule RIPK1 kinase inhibitor. These studies are largely in agreement that RIPK1 kinase inhibition protects β-cell lines as well as mouse, porcine, and human islets from cytotoxicity *in vitro*^36–38^. In addition, Nec-1 blocks β-cell loss in a zebrafish model of overnutrition^39^, and treatment of islets with Nec-1 prior to transplantation in mice improves insulin release and glycemia *in vivo*^40^. Here, we evaluated two recently reported small molecule RIPK1 inhibitors to test their effects on TNFα+IFNγ-induced β-cell death. To that end, we examined a small molecule RIPK1 kinase inhibitor (SZM’679^44^) with beneficial effects in a preclinical model of Alzheimer’s disease and a RIPK1-directed proteolysis targeting chimera (PROTAC, LD4172^45^) that enhances anti-tumor immunity. Like our findings in NIT-1 Ripk1Δ cells, both molecules significantly reduced TNFα+IFNγ-induced NIT-1 cell death over 24 hours. However, only SZM’679, but not LD4172, effectively blocked cytokine-induced caspase 3/7 activation. We postulate this discrepancy is related to loss of RIPK1 kinase activity alone versus loss of both RIPK1 kinase activity and scaffolding function, as previous studies identified different phenotypes related to these domains^25,59–61^. Given that small molecule RIPK1 inhibitors are well-tolerated and under clinical evaluation for treatment of other diseases^62–65^, our findings suggest therapies targeting RIPK1 should be evaluated as agents to protect β-cells in early T1D.

There are some limitations to the current study. These studies utilized an immortalized mouse β-cell line derived from T1D-prone NOD mice. Although this approach allowed us to interrogate β-cell autoimmunity in this widely used preclinical model, additional studies of RIPK1 in primary islets and human β-cells are needed to support translation of our findings. Similarly, we modeled RIPK1 deficiency using two small molecules or CRISPR-Cas9-mediated Ripk1 gene editing. Additional genetic approaches are needed to test RIPK1 kinase activity versus protein expression in models of autoimmune diabetes. To these ends, future studies to evaluate RIPK1 kinase dead (Ripk1^D138N/D138N^) or β-cell-specific RIPK1 knockout mice on the NOD background would be informative. With respect to kinome profiling, kinase binding beads are designed to enrich for kinases in active confirmation, but binding can be affected by activity state, abundance, and affinity of the bead mixture for each kinase. Finally, although our study interrogated the role of RIPK1 in β-cells, RIPK1 also plays important roles in T-cells^66,67^. Thus, studies on the role of T-cell RIPK1 in the pathogenesis of autoimmune diabetes are warranted.

In summary, this study provides novel insights into the mechanisms by which RIPK1 regulates cytokine and autoimmune stress in β-cells. We identified key roles for RIPK1 in T1D-prone β-cell fate, including actions on cytotoxicity-related gene expression, kinase signal transduction, and cell death. Notably, we demonstrated that RIPK1-deficient β-cells are resistant to T-cell mediated autoimmune killing both *in vitro* and *in vivo.* Although RIPK1 is under investigation as a therapeutic target in several diseases including ulcerative colitis^63^ and rheumatoid arthritis^68^, this work sheds new light on the potential of RIPK1-targeting strategies to oppose β-cell loss in the pathogenesis of T1D.

## METHODS

### Cell culture

NIT-1 β-cells derived from NOD/ShiLtJ mice^69^ were cultured in 25 mM glucose DMEM media containing 10% fetal bovine serum (FBS), 1% sodium pyruvate, and 1% penicillin-streptomycin. Ripk1 gene-edited (Ripk1Δ) and control (CTL) NIT-1 β-cell lines were generated as described previously^50^. Briefly, two gRNA sequences targeting exons 2-3 of mouse Ripk1 (5’-GAGAAGACAGACCTAGACAG-3’ and 5’-CCAAATGGTCTGATAGATAT-3’) and a non-targeting control gRNA (5′-TAAAAACGCTGGCGGCCTAG-3′) were cloned into lentiCRISPR v2 vectors (Addgene, #52961), then lentiviral expression of gRNA and Cas9 followed by selection with puromycin was used to establish NIT-1 Ripk1Δ and NIT-1 CTL cell lines. Mutation of the Ripk1 target region was verified by genomic DNA PCR^50^. Mouse islets were cultured in RPMI 1640 media with 11.1 mM glucose, 10% fetal bovine serum (FBS), 1% sodium pyruvate and 1% penicillin-streptomycin. EndoC-βH1 cells were cultured in 5.5 mM glucose DMEM containing 2% fatty acid-free bovine serum albumin (BSA), 50 μM 2-β-mercaptoethanol, 10 mM nicotinamide, 5.5 μg/mL transferrin, 6.7 ng/mL sodium selenite, 4 mM L-glutamine, 1mM pyruvate, 100 U/mL penicillin, and 100 μg/mL streptomycin on plates coated with 1.2% Matrigel containing 3 μg/mL fibronectin. Cells and islets were maintained in a 37°C incubator with 5% CO2. The following reagents were applied to cell cultures: TNFα (40 ng/mL, mouse: CYT-252, Prospec, Rehovot, Israel), IFNγ (100 ng/mL, mouse: CYT-358, Prospec, Rehovot, Isreal), SZM’679^44^ (1 μM, WuXi AppTec, Shanghai, China), LD4172^45^ (100 nM, Dr. Jin Wang, Baylor College of Medicine).

### Quantitative real time polymerase chain reaction (qPCR)

Total RNA from was isolated from cells using the High Pure RNA Isolation Kit (Roche, #11828665001, Basel, Switzerland), then reverse transcribed with the QuantiTect Reverse Transcription Kit (Qiagen, #205311, Venlo, Netherlands), and cDNA was subjected to qPCR. Data were normalized to *Gapdh* RNA levels and expressed as fold relative to control using the ΔΔCT method. All qPCR data points represent means of triplicate technical determinations. Taqman probes (ThermoFisher Scientific, Waltham, MA) were used to quantify mRNA expression of the following targets: *Ripk1* (Mm00436354_m1) and *Gapdh* (Mm99999915_g1).

### Immunoblot analysis

Cell lysates were prepared in buffer containing 50 mM Tris pH 7.5, 2 mM EGTA, 10 mM EDTA, 10 mM NaF, 0.2% Triton X-100 and protease and phosphatase inhibitors (Millipore Sigma, #04693116001 and #0490683700, Burlington, MA). Islets were lysed using a 1% SDS lysis buffer containing 50 mM Tris pH 7.5, 2 mM EGTA, 10 mM EDTA, 10 mM NaF, 150 mM NaCl, 0.2% Triton X-100 and protease and phosphatase inhibitors. Cell lysates were centrifuged at 10,000xg for 10 minutes, then supernatants were collected, and protein concentration was determined by BCA assay (ThermoFisher Scientific, #23227). Equal amounts of total protein were separated on 4-20% SDS-PAGE gels (BioRad, #4561093, Hercules, CA), then transferred to PVDF membranes (Millipore Sigma, #IPVH00010). Proteins were visualized using the following primary antibodies: total RIPK1 (BD Biosciences, #610458, Franklin Lakes, NJ, 1:1000), phospho RIPK1 Ser166 (Cell Signaling, mouse: #31122S, human: #65746, Danvers, MA,, 1:500), β-ACTIN (Abcam, ab8226, Cambridge, UK, 1:2000), GAPDH (Abcam, #ab9485). Primary antibodies were detected with goat anti-mouse 800 (LI-COR, #926-32210, Lincoln, NE, 1:5000) or donkey anti-rabbit 680 (LI-COR, #926-68071, 1:5000) IRDye secondary antibodies, then visualized with a LI-COR CLx imaging system. Representative immunoblot images are shown.

### Immunohistochemistry

Mouse pancreas was extracted and fixed in 10% neutral buffered formalin (ThermoFisher Scientific, # SF100-4). Fixed pancreas samples were embedded in paraffin, processed, and 4 μm sections were cut as described previously^51,70^. Human pancreas sections were provided by the Integrated Islet Distribution Program. Briefly, β-cells were stained using an anti-insulin antibody (Millipore Sigma, #I2018;) followed by goat anti-mouse Alexa Fluor 488 (ThermoFisher Scientific, #A-11001) and an anti-RIPK1 antibody (BD Biosciences, #610458) followed by goat anti-rabbit Alexa Fluor 568 (ThermoFisher Scientific, #A-11011). Sections were then mounted with polyvinyl alcohol and images were captured on a Zeiss LSM710 confocal microscope (Zeiss, Oberkochen, Germany).

### Caspase 3/7 activity assays

Caspase 3/7 activity was quantified using Caspase-Glo 3/7 luminogenic caspase 3/7 substrate (Promega, #G8090, Madison, WI). Briefly, cells were lysed, caspase 3/7 substrate and buffer were added for 1 hour, then luminescence was quantified using a microplate reader (BioTek Synergy H1, Agilent, Santa Clara, CA) and data were reported relative to control treatment conditions for each replicate^50^. Alternatively, caspase 3/7 activity was quantified in single cells in real time using Incucyte Caspase 3/7 Green (Sartorius, #4440, Göttingen, Germany, 5 μM) and live cell imaging and analysis (Incucyte S3, Sartorius). Briefly, caspase 3/7 activity was quantified hourly and reported as caspase 3/7 green positive objects per μm^2^ phase positive cell area for each condition and replicate.

### Cell death assays

Cell death assays were conducted using Sartorius Incucyte S3 live-cell imaging and analysis instruments. Briefly, NIT-1 cells were grown in 48-well plates and cell culture media containing treatments of interest and Sytox Green (ThermoFisher Scientific, #S7020, 100 nM), a membrane impermeable DNA-binding dye that labels dying cells, was added immediately prior to imaging. Four images from each well were collected with a 10X objective each hour, and cell death was quantified as the number of Sytox positive objects at each time point corrected to μm^2^ phase positive cell area for each condition and replicate. In separate experiments, cell culture media containing both Cytotox Red (Sartorius, #4632, 250 nM) and Caspase 3/7 Green (Sartorius, #4440) were applied to label cell death and caspase 3/7 activation in tandem in single cells. Briefly, dyes were added to cell cultures and allowed to equilibrate for 4 hours. Immediately prior to imaging, TNFα and IFNγ were applied, and images were collected hourly for up to 24 hours. Cytotox Red^+^, Caspase 3/7 Green^+^ dual positive objects were considered cells undergoing apoptosis, and Cytotox Red^+^, Caspase 3/7 Green^-^ objects were considered cells undergoing non-apoptotic cell death. Object counts were corrected to μm^2^ phase positive cell area for each condition and replicate. We also colabeled cells with Sytox Green and Nuclight Red (Sartorius, #4717), a membrane permeable DNA binding dye that labels nuclei in all cells. Using this approach, we correlated Sytox Green^+^/μm^2^ phase^+^ area values with Sytox Green^+^ objects/Nuclight Red^+^ objects values within wells to relate our measurements to percent death.

### RNA sequencing and data analysis

Total RNA was extracted from either control (CTL) or Ripk1 deficient (Ripk1Δ) NIT-1 β-cells treated with either vehicle or TNFα+IFNγ. Six biological replicates were generated from each of the four conditions, yielding 24 samples total. RNA-seq libraries were prepared and sequenced on an Illumina NovaSeq 6000 platform (Illumina, San Diego, CA), and ∼30 million paired-end reads (2×100 bp) were generated per sample. Raw sequencing reads were aligned to the mouse reference genome (GRCm39) using STAR aligner (v2.7.11a). Gene-level quantification was performed using featureCounts (subread package v2.0.6), for counting reads mapped to exons of protein-coding genes. Differential expression analysis was conducted using DESeq2 (v1.42.1) in R (v4.3.1). Raw count data were filtered to retain genes with at least 10 reads in a minimum of 3 samples. Normalization and dispersion estimation were performed using DESeq2’s median-of-ratios method and empirical Bayes shrinkage, respectively. Statistical significance was determined using the Wald test with Benjamini-Hochberg correction for multiple testing (FDR<0.05). The differential expression analysis utilized accounts for genotype alone (CTL vs Ripk1Δ), treatment alone (vehicle vs TNFα+IFNγ), and the interaction between genotype and treatment. Two ranking approaches were used to highlight distinct aspects of the differential expression results. For volcano plots, genes were ranked by statistical significance alone, with the top 10 most significantly upregulated and downregulated genes (lowest FDR values) labeled for each comparison. For heatmap visualizations, a combined ranking approach that integrates statistical significance and fold change magnitude was employed. Briefly, for each comparison, Kruskal-Wallis tests were used to calculate p-values for differential expression between groups, with subsequent Benjamini-Hochberg correction to obtain false discovery rates (FDR). Log2 fold changes were calculated as the log2 ratio of mean expression between comparison groups. Genes were then ranked separately for a) FDR values in ascending order, with genes having lower FDR values receiving better ranks, and b) magnitude of log2 fold change in descending order, with genes having larger absolute fold changes receiving better ranks. These ranks were then summed to form a combined rank score, with lower combined rank scores indicating genes with greater statistical robustness and magnitude of fold change. The top 30 genes in this combined ranking were selected for heatmap visualization. Separately, reads for transcripts of interest in curated gene sets were subjected to heatmap visualization.

### Multiplex kinase inhibitor bead affinity chromatography-mass spectrometry (kinome profiling)

For multiplex kinase inhibitor bead affinity chromatography-mass spectrometry (MIB-MS, kinome profiling), NIT-1 cells were treated, washed in cold PBS, harvested, spun, supernatants removed, then cell pellets were flash frozen and stored at -80°C for subsequent analysis. Frozen cell pellets were lysed in buffer containing 50 mM HEPES, 150 mM NaCl, 0.5% Triton X-100, 1 mM EDTA, and 1 mM EGTA, 10 mM NaF, 2.5 mM sodium orthovanadate, complete protease inhibitor cocktail (Millipore Sigma, #04693116001), and phosphatase inhibitor cocktails II and III (Millipore Sigma, #524636 and # 524631), then samples were sonicated and cleared lysates were equalized at 0.76 mg total protein per replicate and brought to 1M NaCl. Lysates were flowed over kinase inhibitor bead resin containing 250 μL of a 50% slurry of seven kinase inhibitors (CTX, VI-16832, PP58, Purvalanol B, UNC0064-79, UNC0064-12, and BKM-120) covalently linked to ECH-Sepharose 4B beads as described previously^53^. Bound kinases were washed with high salt (1M NaCl), low salt (150 mM NaCl), and buffer (50 mM HEPES, 0.5% Triton X-100, 1 mM EDTA, 1 mM EGTA, pH 7.5), eluted with boiling 0.5% SDS and 1% β-mercaptoethanol in 100 mM Tris-HCl, treated with DTT and Iodoacetamide, spin-concentrated to 100μL (Millipore Sigma, #UFC8010D), and subjected to methanol-chloroform precipitation. Proteins were digested with trypsin overnight at 37°C, extracted with water-saturated ethyl acetate, dried in a speed-vac, desalted with C-18 spin columns (Agilent Technologies, #5188-2750, Santa Clara, CA), and resuspended in 0.1% formic acid. Mass spectrometry was performed on a Vanquish Neo UHPLC coupled to an Orbitrap Exploris 480 (both ThermoFisher Scientific). Resulting RAW files were processed using an *in silico* DIA-NN 2.0 predicted spectral library^71^ generated from the UniProt/Swiss-Prot mouse database. LFQ intensities from unique genes for all annotated kinases with at least two unique peptides were analyzed in R (v.4.5). Relative intensities were log2 transformed, filtered to include annotated kinases with at least three valid values in a treatment group, and missing values imputed from the normal distribution for each column using a Perseus-style imputation^72^. Two sample, unpaired Student t-tests of log2LFQ intensities were performed and plotted using ggpubr (p<0.05, log2FC <|1| significance cut-off). The z-score of each kinase was calculated, and the Euclidian distance was plotted using the pheatmap R package.

### Islet and splenocyte isolations

Mice used in this study were maintained with ad libitum access to food and water under protocols approved by the Indiana University (IU) Institutional Animal Care and Use Committee. Islets were isolated by the IU Center for Diabetes and Metabolic Diseases (CDMD) Islet and Physiology Core and maintained in RPMI 1640 media with 10% fetal bovine serum (FBS), 1% sodium pyruvate, and 1% penicillin-streptomycin. Diabetic splenocytes were harvested from spontaneously diabetic female NOD mice as previously described^73^. Briefly, spleens were mechanically dissociated and passed through a 70 μm cell strainer to obtain a single-cell suspension. Red blood cells were lysed using a hypotonic buffer (Millipore Sigma, #11814389001), and leukocytes were subsequently washed with PBS and counted with an automated cell counter (BioRad, #1450102).

### ELISPOT assay

NIT-1 cells were seeded in 6-well plates at a density of 2×10⁶ cells per well and grown overnight. The following day, cells were treated with 5 μM thapsigargin (Millipore Sigma, #T9033) for 1 hour. After treatment, cells were enzymatically detached, resuspended in DMEM with 10% FBS and 2×10⁴ NIT-1 cells per well were added to pre-coated and blocked 96-well murine IFNγ ELISPOT plates (BD Biosciences, #551083). Splenocytes isolated from a spontaneously diabetic female NOD mouse were added at 7×10⁵ cells per well. Phytohemagglutinin (PHA, 2 μg/mL) was used as positive controls, and wells with splenocytes but no NIT-1 cells served as negative controls. NIT-1 cells and splenocytes were co-cultured (1:35 ratio) for 24 hours at 37 °C in 5% CO₂. After incubation, cells were removed, and plates were washed with PBS containing 0.1% Tween-20. Spot quantification and analysis were conducted using the ImmunoSpot S6 Universal-V Cell Counter Analyzer (Cellular Technology Limited, Shaker Heights, OH).

### *In vitro* NIT-1 β-cell and diabetic splenocyte co-culture

NIT-1 cells were labeled with CellTrace Violet (ThermoFisher Scientific, # C34557) and seeded at 1×10⁵ cells per well in 48-well plates. The following day, cells were treated with 5 μM thapsigargin for 1 hour, washed with PBS to remove residual compound, and co-cultured with freshly isolated splenocytes from a diabetic NOD mouse at an effector-to-target (E:T) ratio of 2:1 for 48 hours. For assessment of T-cell activation, cells were stained with fluorescently conjugated anti-CD8 (BioLegend, #100712, San Diego, CA) and anti-CD69 (BioLegend, #104512) antibodies and analyzed by flow cytometry. NIT-1 cells were identified by CellTrace Violet gating and assessed for survival using Zombie NIR viability dye (BioLegend, 423105). All samples were acquired on a Cytek Aurora flow cytometer (Cytek Biosciences, Fremont, CA) and analyzed using FlowJo software (version 10.1.1).

### *In vivo* autoimmune killing and NIT-1 β-cell bioluminescence imaging

NIT-1 CTL and Ripk1Δ cell lines were engineered to constitutively express firefly luciferase (Luc2) under the control of the EF1α promoter via lentiviral transduction. A total of 1 × 10⁷ control (CTL) and Ripk1-deficient (Ripk1Δ) luciferase-expressing NIT-1 cells were implanted subcutaneously on opposite flanks of immunodeficient NOD.scid.gamma (NSG) mice. β-cell autoimmunity was induced by intravenous injection of 1×10⁷ splenocytes freshly isolated from spontaneously diabetic female NOD mice. For bioluminescence imaging, D-luciferin (Gold Biotechnology, #LUCK-100, St. Louis, MO) was administered intraperitoneally at 150 mg/kg, then graft luminescent signals were acquired using the IVIS Spectrum Imaging System (PerkinElmer, Waltham, MA) 1, 2, 4, 7, 12, 14, and 19 days post implant.

### Statistical Analysis

Statistical tests were conducted with GraphPad Prism 10 software (GraphPad, San Diego, CA). Unpaired, two-tailed Student’s t-tests were used to analyze data sets with two groups, and one-way analysis of variance (ANOVA) tests were used to analyze data sets with more than two groups. Significant ANOVA results were followed with Holm-Sidák post-tests to analyze differences between groups of interest. Paired statistics were applied when all biological replicates contained all experimental conditions and data was collected and analysed at the same time within a replicate. Simple linear regression was used to determine the correlation between two measurements of interest. Data are presented as mean ± standard error with a value of p<0.05 considered significant.

## DATA AVAILABILITY

All datasets are available from the corresponding author on reasonable request.

## AUTHOR CONTRIBUTIONS

C.J.C.: Conceptualization, formal analysis, investigation, methodology, writing- original draft, writing- review and editing; N.M., A.H.K., E.G.M., N.A., C.D., C.A.B., M.P.: Investigation, formal analysis, methodology, writing- original draft, writing- review and editing; D.K., K.A.C., A.S.M., R.C.S.B.: Investigation, writing- review and editing; T.S.J., S.P.A., S.Y., E.P.C.: Conceptualization, methodology, funding acquisition, writing- review and editing; A.T.T.: Conceptualization, data curation, formal analysis, funding acquisition, investigation, methodology, supervision, writing- original draft, writing- review and editing.

## ADDITIONAL INFORMATION

**Correspondance** should be addressed to Andrew T. Templin or Christopher J. Contreras.

### Acknowledgements

We thank Dr. Jin Wang (Baylor College of Medicine) for providing LD4172, a small molecule proteolysis targeting chimera (PROTAC) used for RIPK1 protein degradation studies. We thank Emily C. Sims (Indiana University School of Medicine Cell Response Technologies Core) for expert assistance with Incucyte data collection and analysis. We thank Kara Orr and Lata Udari (IU Center for Diabetes and Metabolic Disease Islet and Physiology Core) for expert assistance with islet isolation. We thank the IU Center for Medical Genomics for expert assistance with RNAseq sample preparation and data acquisition. We acknowledge BioRender for providing graphic design software used to create figure schematics. Some data from this study were presented at the American Diabetes Association 83^rd^ and 85^th^ Scientific Sessions.

